# Distributional Data Analysis Uncovers Hundreds of Novel and Heritable Phenomic Features from Temporal Cotton and Maize Drone Imagery

**DOI:** 10.1101/2025.09.05.674557

**Authors:** Aaron J. DeSalvio, Marcos Matabuena, Alper Adak, Mustafa A. Arik, Serina M. DeSalvio, Seth C. Murray, Raymond K.W. Wong, Jode Edwards, Natalia de Leon, Shawn M. Kaeppler, Dayane C. Lima, Candice N. Hirsch, Addie Thompson, David M. Stelly

## Abstract

Genomic and phenomic analyses suggest additional heritable phenomic features can improve modeling of important end traits like senescence or yield. Field phenotyping generally uses trait values averaged across individual experimental units (plants or numerous plants within plots), ignoring the full distributional pattern of collected measures. Images of plants or plots, as captured by drones (unoccupied aerial vehicles / UAVs / drones), can be viewed as individual distribution functions that capture biological information. This study introduces and validates distributional data analysis in two crops and experiment types – cotton (*Gossypium hirsutum* L.) single plant vegetation index (VI) analysis and maize (*Zea mays* L.) plot-level yield predictions. In both crops, the concept of within-day variance decomposition was demonstrated. In cotton, genotypes exerted significant influences on temporal quantile functions of VIs. Maize yield prediction using distributional data with elastic-net regression indicated improvements in yield prediction between 12.7%-21.6% with quantiles outside the conventionally used median responsible for added predictive power. A novel data visualization method for per-pixel heritability allowed distributional features to be explainable and interpretable. These results have implications for future plant phenomic studies, indicating that distributional data analysis applied across temporal imagery captures novel, heritable, and interpretable biological signal that is lost when working with conventional measures of central tendency such as mean or median summary values of experimental units.

**Significance:** Repeated aerial imaging of agricultural experiments produces image data sets that capture plant development in high spatial and temporal resolutions. Frequently, images are summarized by measures of central tendency, such as mean or median values. Here, functional data distributional methods were applied to cotton (*Gossypium hirsutum* L.) and maize (*Zea mays* L.) image data, capturing more information than standard approaches. Cotton genotypes significantly impacted distributional spectral data while in maize, distributional data enabled more accurate predictions of grain yield versus models trained with median data alone. Distributional data were more explainable by genetics, with novel data visualization techniques able to shine light on specific parts of plant imagery with high and low genetic variance.

## Background

### Plant Phenomics

Phenomics is defined as the “acquisition of high-dimensional phenotypic data on an organism-wide scale” (Houle et al., 2010). Phenomics has witnessed rapid expansion since it was first reviewed in a plant-specific context (Furbank & Tester, 2011) owing to advancements in imaging and complementary molecular genetic technologies, and has been further reviewed to show value in biological predictions and discoveries (Kumar et al., 2015; Lyu et al., 2017; Tardieu et al., 2017; White et al., 2012; Yang et al., 2020; Zavafer et al., 2023; Zhao et al., 2019). Growing interest is similarly occurring across humans, animals, and microbes (Acin-Albiac et al., 2020; Pérez-Enciso & Steibel, 2021; Ying, 2023). Zavafer et al. (2023) define phenomics as “the discipline of biology that studies a hypervolume” (the phenome), comprising every phenotype a genome (G) can express across environments (E), defined as a location/year combination, and time (T). By interrogating that G×T×E hypervolume, phenomics links genotype to function and plasticity of functions. Temporal measurements reveal when genes act, environmental gradients expose how they respond to stress or optimal conditions and together uncover mechanisms that shape fitness and adaptation. Because G×T×E effects dictate cumulative end traits of importance, plant phenomic data sets have been exploited in numerous ways, such as for the analysis of leaf senescence in a greenhouse setting (Lyu et al., 2017), prediction of grain yield in maize (Adak et al., 2021), wheat (*Triticum aestivum* L.) (Jackson et al., 2023; Winn et al., 2023), chickpea (*Cicer arietinum* L.) and dry pea (*Pisum sativum* L.) (Zhang et al., 2021), coffee (*Coffea canephora*) (Adunola et al., 2024), soybean (*Glycine max* (L.) Merr.) (Parmley et al., 2019), among others. These studies show improved discovery and end trait prediction using high-heritability features captured through time.

Contemporary studies capture phenomic data using technologies such as near infrared spectroscopy (NIRS) (Lane et al., 2020; Rincent et al., 2018; Robert et al., 2022; Zhu et al., 2021), in-field robots (Mueller-Sim et al., 2017; Underwood et al., 2017; Virlet et al., 2016), greenhouse imaging systems (Kaya, 2025), or through use of unoccupied aerial vehicles (UAV/UAS/drones), as reviewed in Guo et al. (2021) and Gano et al. (2024). Drone imaging offers distinct advantages of scalability and portability while enabling capture of dense temporal and spectral data at high resolution (0.1cm – 1.0cm/pixel) compared to satellite imagery (30cm – 10m/pixel) or cameras mounted on planes. Images stitched into orthomosaics and used to build 3D point clouds enable temporal spectral and structural feature extraction at single-plant, plot, or whole-field levels. Open source tools such as *FIELDimageR* (Matias et al., 2020; Pawar & Matias, 2023) and *FIELDimagePy* (Chatterjee et al., 2025) provide phenomic feature extraction interfaces for mean and median vegetation index (VI) values. The resulting VI values are ratios of bands captured by a sensor within boundaries of individual experimental units (plants or plots), which tend to be more robust and more predictive than raw spectral bands. However, a wealth of untapped phenomic information exists in readily available distributional data from images that has not yet been studied in plants.

### Functional and Distributional Data Analysis

Across many facets of biology, routine (scalar) summaries of central tendency (e.g., mean/median VI values of plant or plot images) incompletely explain differences in end traits, with Anscombe’s quartet and the Datasaurus dozen serving as visual representations of this phenomenon. At single-plant resolution, naturally occurring senescence often begins from the edges of a leaf at its base, while localized leaf senescence can occur in the event of unevenly applied environmental stress (Lim et al., 2007). Plant-to-plant variations could be overlooked when routine summary statistics are used for each plant. At the plot level, factors such as variable germination progression (Xue et al., 2021) or seedling uniformity (Muldoon & Daynard, 1981) can explain differences in terminal measures. Scalar summaries that aggregate image data are inadequate to parse these subtle phenomena, especially when these summaries are captured temporally.

Functional data analysis (FDA) (Crainiceanu et al., 2024; Ramsay & Silverman, 2005; Wang et al., 2016), is a statistics discipline that treats continuous data (e.g., temporal) as individual data objects for each experimental unit. It is used in medical functional magnetic resonance imaging (fMRI) (Huang et al., 2017), which benefits from uniform collection formats over short time windows. Extension of these analytic methods to entire organismal life spans is complicated by irregular and sparse measurement intervals across disparate environments. Computational efficiency requirements for large data sets, violations of error independence assumptions, and nongenetic correlations of neighboring time series data (e.g., field spatial variation of nearby plants) are challenges similarly faced in applying functional data analysis directly to both brain imaging studies (Tian, 2010) and agricultural data sets.

Human studies have exploited the richness of distributions while avoiding information loss inevitable when relying on scalar summaries. Distributional data analysis of wearable accelerometers has provided benchmark data, enabling studies that fall within the scope of digital phenotyping (Onnela, 2021). Distributional data analysis of biosensor data was first used in continuous glucose monitoring (CGM) (Matabuena et al., 2021), with multivariate representations of CGM data outperforming classical non-distributional CGM indices (Matabuena et al., 2024). Accelerometry data has enabled modeling cumulative distribution functions for individual subjects (Matabuena & Petersen, 2023), implementing subject-specific quantile functions for studying gait and Alzheimer’s Disease (Ghosal et al., 2023) and uncovering links with cognitive evaluation scores (Ghosal & Matabuena, 2024). Other examples include analysis of single-cell data (Tiberi et al., 2023), causal effect estimation (Wang et al., 2023), quantile regression (Waldmann, 2018), and distributional regression (Klein, 2024).

In most plant studies, the replication of experimental units both within and across environments over time allows heritability metrics to objectively demonstrate value in measurements. Here, explainable biological insights were made from multispectral drone image data collected over individual cotton (*G. hirsutum* L.) plants and RGB (red, green, blue) image data from maize (*Zea mays* L.) hybrid plots, differentiating these methods from deep learning approaches, which suffer from a lack of or indirect explainability. Distributional information from temporal plant images uncovered novel phenomic features with heritability superior to conventional mean-and median-based summaries of phenomic data. Genetic variance and heritability of VI distributional data in cotton and maize, along with grain yield prediction followed by a feature importance analysis in maize using elastic-net regression, serve as objective metrics of novel value from distributional and FDA methods across two crop species and multiple environments.

## Methods Overview

Graphical summary of data collection and extraction methods of individual cotton plants is presented in Fig. 1. All code referenced in the Methods section is available in GitHub [https://github.com/ajdesalvio/phenomic-distributional].

**Figure 1.**
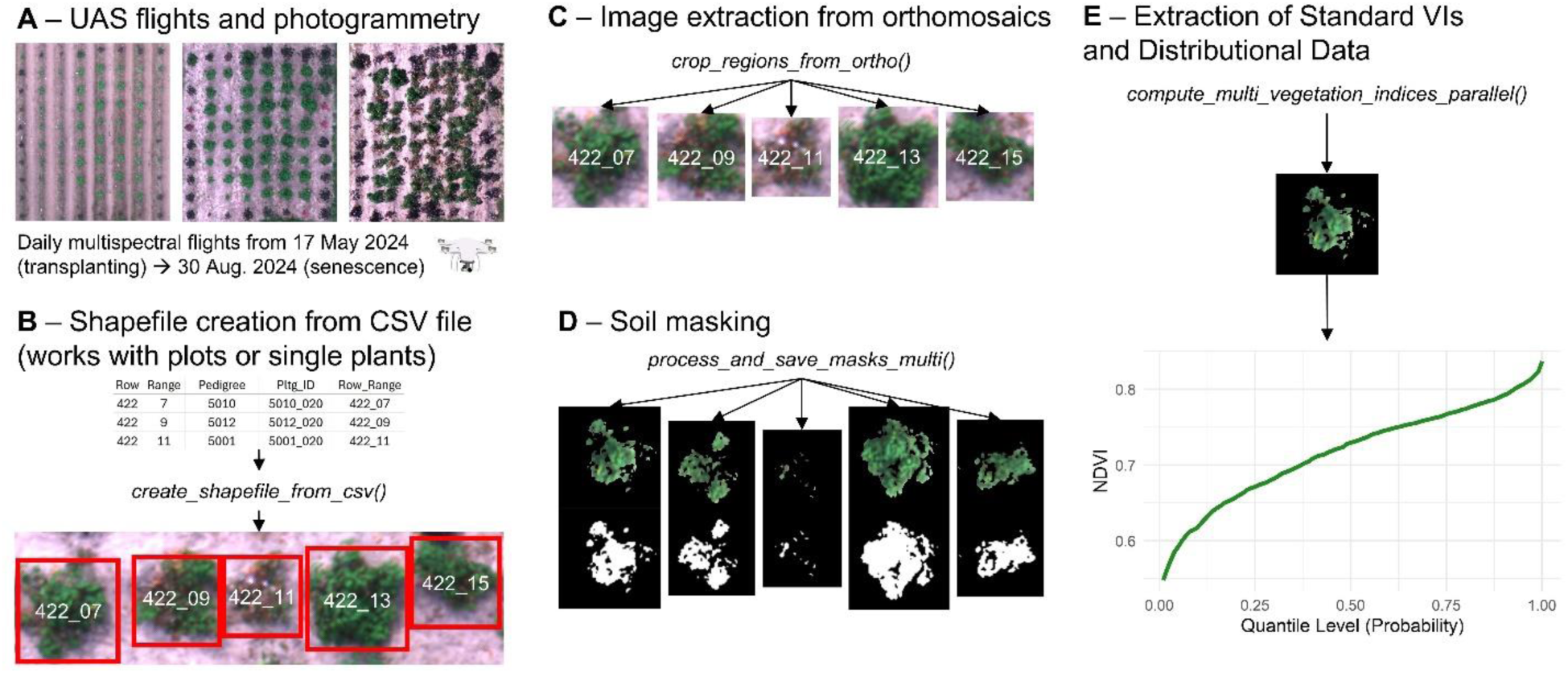
Graphical summary of data collection and extraction methods of individual cotton plants. (**A**) Multispectral drone flights for cotton were conducted daily, and images were stitched into orthomosaics. (**B**) Shapefiles were created with bounding boxes overlaid on top of single plants; (**C**) Single-plant images were extracted from each orthomosaic; (**D**) Soil was masked from each single-plant image; (**E**) Standard vegetation indices (mean and median values per plant) as well as VI distributional data were calculated, with the quantile function for a single plant at one time point shown as an example. Italicized text indicates custom Python functions.

## Cotton Methods

### Field Experiment Design

A field experiment in College Station, TX, between April and August 2024 was conducted on upland cotton (*G. hirsutum*) BC_5_ backcross-inbred lines containing introgression segments from wild Hawaiian cotton *G. tomentosum* (Nutt. Ex Seem.) detailed in DeSalvio et al. (2024). Genotypes included 3 chromosome substitution lines (CSLs), 12 chromosome segment substitution lines (CSSLs), and a control genotype, Texas Marker 1 (TM-1; Supplementary Table 1; Table S1), totaling 16 unique genotypes. Greenhouse germinated seeds were mechanically transplanted on 12 April 2024 into a randomized complete block design (RCBD) with 10 blocks. Each block had 48 spaced plants, with three replicates of each genotype, totaling 480 plants, with 479 available for analysis after one succumbed to transplantation shock. Further details regarding the field layout are provided in Supplementary Methods 1 (Methods S1).

### Cotton Ultra-Dense Temporal Phenomic Data and Photogrammetry

Daily cotton drone flights began 17 April 2024 and ended 30 August 2024. Of 106 completed flights, 101 were suitable for downstream analysis. Flights at 46, 71, 72, 99, and 100 days after transplanting (DAT) were removed due to quality issues. Five-band multispectral images were captured with a DJI Phantom 4 Multispectral drone. Sensor details and photogrammetry methods are described in Methods S1.

### Mixed Function-On-Scalar Distributional Regression Model for Phenomic Data

The model from Matabuena and Crainiceanu (2024) facilitated testing the effect of Genotype, a functional fixed effect (Loewinger et al., 2025) on temporal quantile functions in cotton. Differences in phenotypic variation are attributable to unique introgressed *G. tomentosum* segments operating within a *G. hirsutum* background, making Genotype the fixed effect of primary interest. Temporal senescence or stay-green phenotypes using mean VI values (DeSalvio et al., 2024) served as the hypothesized behavior of the CSLs and CSSLs (Table S1). The implementation of the model starting from orthomosaic image files is summarized as follows: i) extract individual experimental units (EUs, single plants in the cotton analysis) at each time point from orthomosaics (Methods S1); ii) using RGB band values, evaluate VIs for every pixel in the EU; iii) calculate a quantile function for the EU; iv) evaluate the quantile function at 100 steps from 0.01,…,1.00; v) aggregate data across time to enable testing for Genotype effects along the quantile continuum using mixed linear models at each quantile level. Random effect covariates, including Range (Y spatial field axis), Row (X spatial field axis), and Replicate (of Genotype) mitigated field spatial variation. A random intercept was specified for each Replicate, while Range and Row were allowed random slopes. The *fastFMM* R package (Loewinger et al., 2025) fit the model, implementing the fast univariate inference (FUI) approach for longitudinal functional data (Cui et al., 2022). The functional mixed model enables assessment of both extent and location within the quantile domain where Genotype exerts a statistically significant impact (if any) on each chosen VI, affording unique insights not offered by conventional mixed models with explainability, unlike many deep learning frameworks.

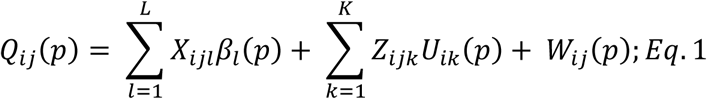

In this model, 𝑄_𝑖𝑗_(𝑝), 𝑝 ∈ [0,1] indicates the random quantile function for plant 𝑖 = 1,…, 𝑛, where 𝑛 = 479, measured at DAT 𝑗 = 1,…, 𝐽_𝑖_. The functional domain is the space of quantile functions, ranging from [0, 1] at intervals of 0.01. All plants had the same number of observations, i.e., 𝐽_𝑖_ = 𝐽 for all plants, with 𝑗 = 101 drone flights. 𝑋_𝑖𝑗𝑙_indicates 𝐿 fixed effects covariates (Genotype was the only fixed effect, containing 16 levels), 𝛽_𝑙_(𝑝) indicates the fixed effect parameter for the Genotype at quantile level 𝑝, 𝑍_𝑖𝑗𝑘_indicates 𝐾 random effects covariates (three: Range, Row, and Replicate), 𝑈_𝑖𝑘_(𝑝) denotes plant 𝑖’s random functional effect at probability 𝑝, and 𝑊_𝑖𝑗_(𝑝) accounts for unexplained residual variation. 𝑈_𝑖𝑘_(𝑝) and 𝑊_𝑖𝑗_(𝑝) are assumed to be zero mean square integrable processes. 𝑊_𝑖𝑗_(𝑝) is assumed to be uncorrelated with the random functional effects 𝑈_𝑖𝑘_(𝑝), however 𝑈_𝑖𝑘_(𝑝) can demonstrate intra-function correlation. Specific algorithmic details regarding fitting Eq. 1 are presented in Matabuena and Crainiceanu (2024).

## Maize Methods

### Field Experiment Design - Maize

The 2020-2021 Genomes to Fields (G2F) maize hybrid experiment phenotypic, environmental, and soil data were described in Lima et al. (2023). Briefly, 1,184 hybrids with publicly available genotyping data were evaluated in 30 locations. Hybrids were produced by crossing doubled-haploid inbreds from the Wisconsin Stiff Stalk MAGIC population with three ex-PVP inbred testers (PHP02, PHK76, and PHZ51) adapted to Northern, Midwest, and Southern locations, respectively. Data presented are from IAH4.2021, MIH1.2020, MNH1.2020, MNH1.2021, TXH1.2020, TXH2.2021, WIH1.2021, WIH2.2020, WIH2.2021, WIH3.2020, and WIH3.2021 environments (combination of location and year), with the first two letters being the state. In each environment, hybrids were planted in a modified RCBD generally with two replicates.

### Maize G2F Temporal Phenomic Data Capture

Maize image capture from the G2F experiments was conducted at different sparse dates in each environment ranging from 10-25 flights in each environment (Supplementary File 1), but the entire developmental trajectory was captured from emergence through senescence in each case. RGB images were used for all presented G2F maize environments. Details for extracting single experimental units (cotton - single plants; maize – plots) with shapefiles and subsequent vegetation index extraction are provided in Methods S1.

### Yield Prediction and Feature Importance Quantification with Maize Distributional Data

To empirically determine whether distributional data could improve predicting and understanding an end trait (e.g., grain yield), the *caret* (Kuhn, 2008) and *glmnet* (Friedman et al., 2021) R packages were used to implement elastic-net regularization (Zou & Hastie, 2005), which was chosen due to the 𝑛 ≫ 𝑝 data structure arising from a modest number of maize hybrid genotypes with a large number of distributional predictors (≤100 quantiles × the number of flights for each environment). The implementation details of elastic-net regression are provided in Methods S2. Variable importance scores and correlations between actual and predicted grain yield values were extracted from the top-performing models. A sensitivity analysis used varying degrees of data truncation after removing extreme quantiles to identify stably important variables for predicting grain yield. Analyses were conducted separately for each of the 11 G2F environments using grain yield (t/ha) best linear unbiased estimators (BLUEs) as response values and distributional VI BLUEs as predictors. BLUEs in both cases were calculated using an analysis of variance model (ANOVA) (labeled AM3; detailed below) after setting the Genotype factor as a fixed effect in *lme4*. BLUEs were best suited for this two-stage analysis (Holland & Piepho, 2024).

### Correlation Analysis of Maize Quantile Phenomic Data

To evaluate redundancy or novelty of temporal quantile data, Pearson correlation matrices were calculated for each DAP within each G2F environment. For within-environment temporal correlation figures, each correlation matrix was stacked by DAP to create a 3D array (100 quantiles × 100 quantiles × number of flights for that environment). The Fisher transformation (Fisher, 1915; Hotelling, 1953) was applied (the *atanh()* R function) element-wise, *z* values were averaged across the flight dimension, and the inverse was obtained with the *tanh()* R function to produce an averaged correlation matrix. For an across-environment average correlation matrix, the array was created by stacking correlation matrices from all environment/DAP combinations before applying the transformation.

### Visualization of Raw Data

The 3-dimensional (3D) nature of VI quantile data is shown in Fig. 2 and accompanying Supplementary Video 1 (Video S1). Normalized difference vegetation index (NDVI), commonly used in plant biology as a proxy for vigor, using a single plant (Genotype 5016, TM-1 replicate 7) tracked across the entire experiment serves as an example. Since quantile functions are monotonic, the NDVI value corresponding to probability 𝑝 + 0.01 cannot be smaller than quantile level 𝑝. For example, at DAT 35 and 𝑝 = 0.20, the quantile function returns the NDVI value at which 20% of the data fall at or below that value (0.58). Fig. 2 reveals substantial ignored variability when relying on only mean or median values. Fig. 2B and 2C demonstrate that data from a single plant can be viewed as a surface existing in 3D space, and 2D emphasizes the total variability in the data for one plant.

**Figure 2.**
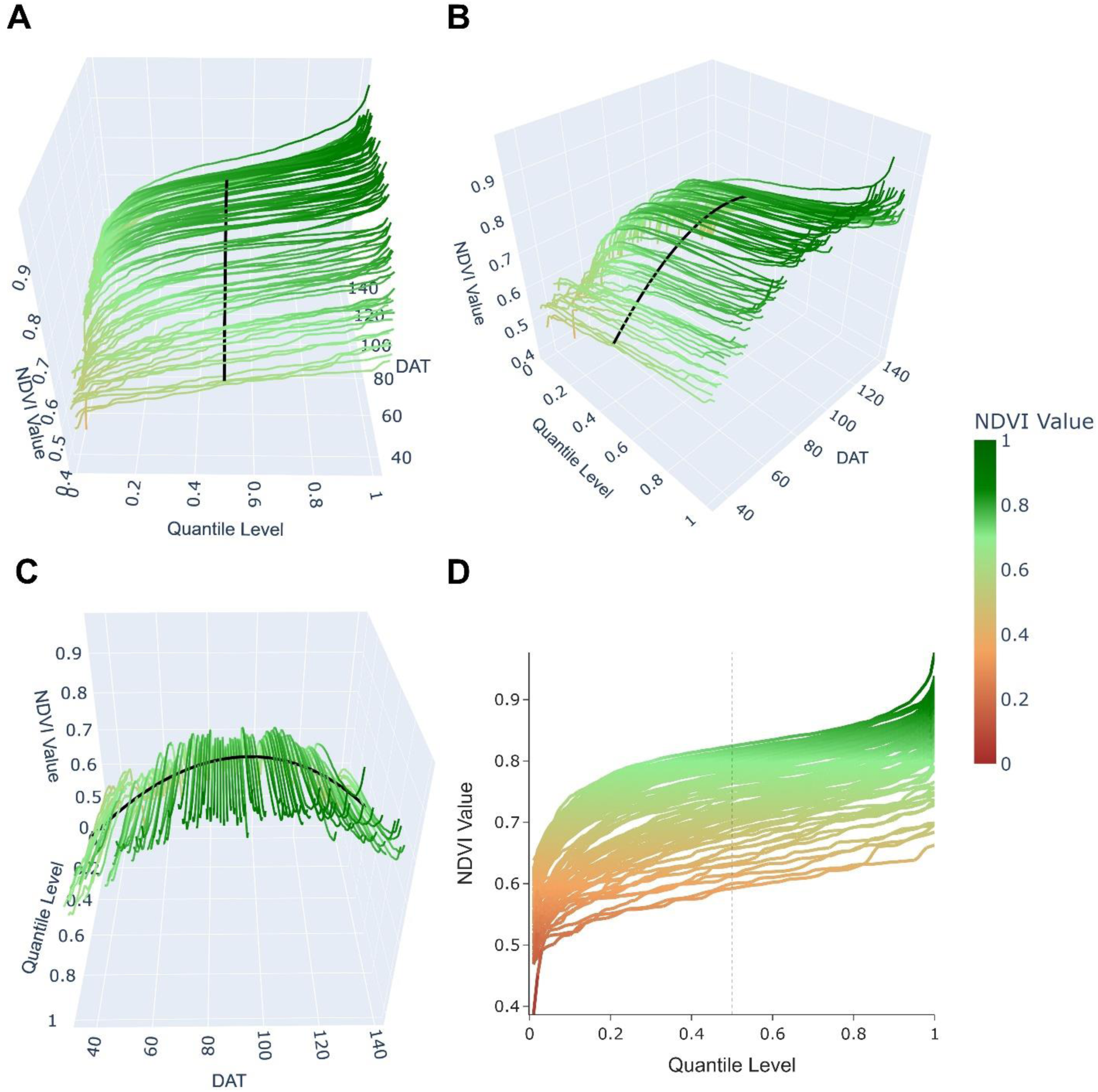
Normalized difference vegetation index (NDVI) quantile functions calculated for a single cotton plant within each drone flight (indexed by days after transplanting, DAT). (**A**) Presents the quantile functional domain of plant 5016_07 (Genotype 5016, TM-1, replicate 7) closest to the reader with a conventional median value for each DAT shown in black at quantile 0.50 (the median trajectory was smoothed with a 1D spline). Quantiles were calculated at each 0.01 interval from 0.01 to 1.00 and plotted as continuous lines; (**B**) intermediate view highlighting that each quantile function’s results are indexed by DAT; (**C**) DAT axis closest to reader showing overall shape of the NDVI quantile function surface across time from transplanting through senescence; (**D**) 2D representation of the temporal quantile data representing the variation in NDVI values among the quantile functions indexed by DAT. Dashed vertical line indicates the median.

## Combined Methods (Cotton and Maize)

### Calculation and Visualization of Heritability

Heritability, a summary statistic of variance components, quantifies the variation attributable to genetics as opposed to other nuisance factors and is a gold standard for objectively comparing different traits or evaluating new measures versus old ones within experiments. Heritability of VI temporal distributional data is detailed in Table 1. All models (Methods S2) were implemented in R version 4.4.2 (R Core Team, 2024) using *lme4* (Bates et al., 2015). Excluding intercepts, all model components were treated as random effects to study the proportion of variation (variance components) each explained within the vegetation index data.

**Table 1.**
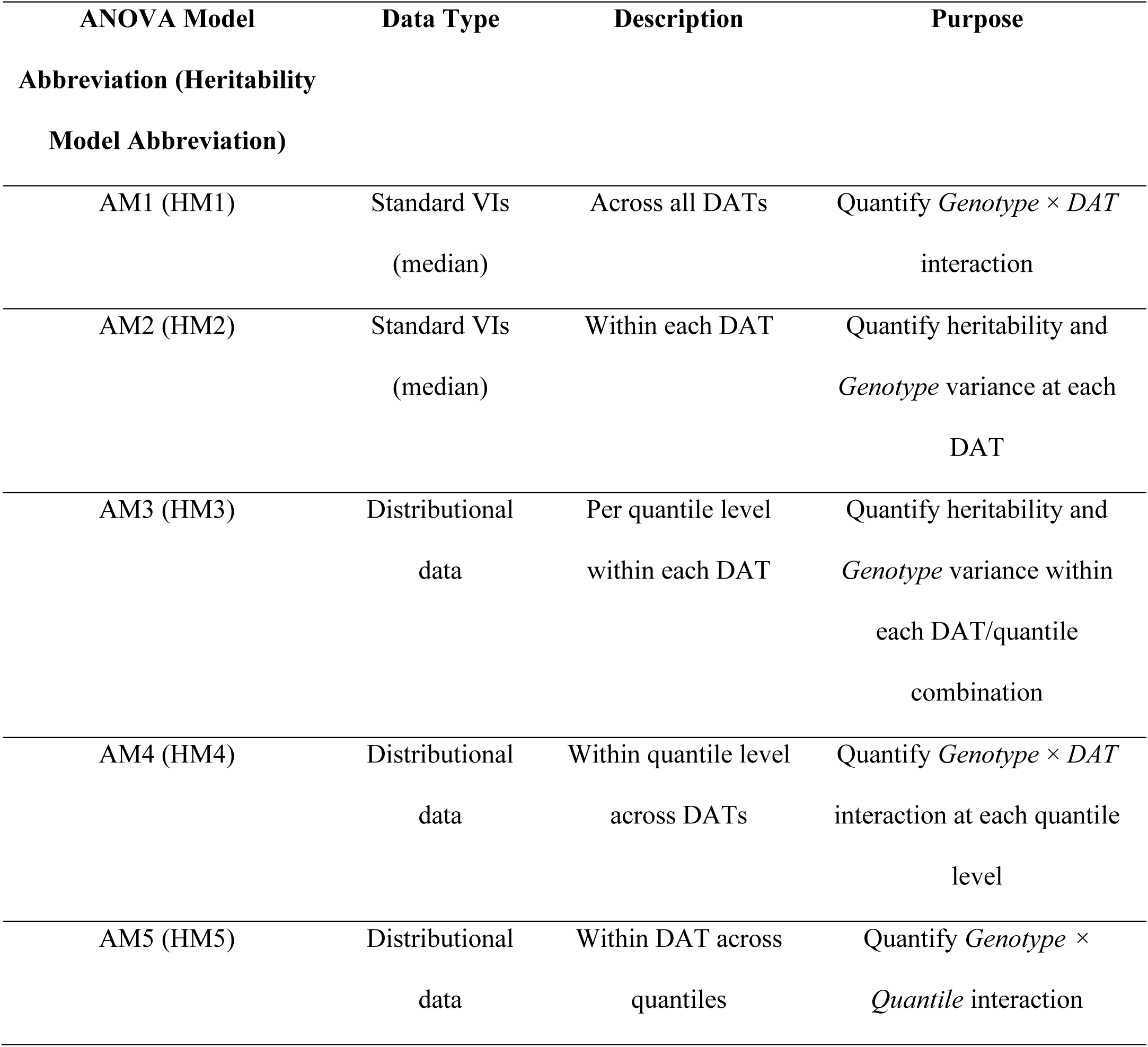
Analysis of variance (ANOVA) models used in this study with model abbreviations and data sets. AM indicates ANOVA model; HM indicates heritability model; DAT - days after transplanting for cotton; DAP - days after planting for maize G2F. Italics indicate the variance component of primary interest.

The distributional nature of images can be visualized as a relation between each pixel’s VI value and the global distribution of VI values from all pixels across all images within a specific flight (since AM3 and HM3 were evaluated at every DAT/quantile level combination). Novel heritability activation maps (HAMs) were created using the empirical cumulative distribution function (eCDF) within each flight date to map each pixel’s VI value to a heritability value obtained from HM3. The eCDF for a set of observations 𝑥_1_, 𝑥_2_,…, 𝑥_𝑛_ is defined as:

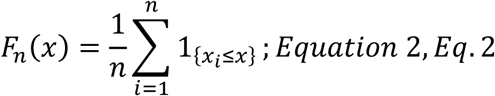

Each 𝑥 is the value obtained after evaluating a pixel’s VI value within one image in one flight date (DAT) and 𝑛 is the total number of valid pixels across all images within that flight. Previously, the VI values at specific quantiles for each image were obtained using *np.quantile()* before analyzing these data with ANOVA models (Table 1; Methods S2). With heritability results from HM3, the eCDF was used to map each pixel’s VI value to its position within the overall distribution of VI values for each flight date and return a quantile. The *input* is a VI value that produces a quantile level, whereas previously the quantile function for an image was evaluated to *output* a VI value at that quantile. For example, if a pixel’s eCDF value within a single-plant image at DAT 86 is 0.60 (the 60^th^ percentile), the heritability value associated with the 60^th^ quantile can be obtained from the ANOVA results (HM3), and a heritability intensity can be assigned to that pixel. Programmatic details of the eCDF method are provided in Methods S2.

## Results

### Function-on-Scalar Regression Quantifies Extent and Location of Cotton Introgression Segments Impacting the Phenome

The function-on-scalar distributional regression model (Eq. 1) allowed significance testing of 16 cotton genotypes to determine whether differences existed in vegetation index (VI) values between the genotypes at each quantile level. The estimated functional coefficients (Fig. 3) are depicted for the ExG2 VI. Interpretation of the functional coefficients is in context that higher ExG2 values correspond to healthier plant tissue (ranging from-1, unhealthy, to 2, healthy; Table 2) and that Genotype 5016 (TM-1 commercial check) was the reference factor level (intercept). Other genotype plots show mean changes from the reference at each quantile level 𝑝. Eq. 1 fit data from all time points, so conclusions about statistical significance can be made for regions of the quantile domain, not the time domain. For quantiles corresponding to 𝑝 > 0.2 up to approximately 𝑝 = 0.95, effects of Genotype 5001 had significantly lower ExG2 values, consistent with its previous senescence phenotype (Supplementary Results; Table S1).

**Figure 3.**
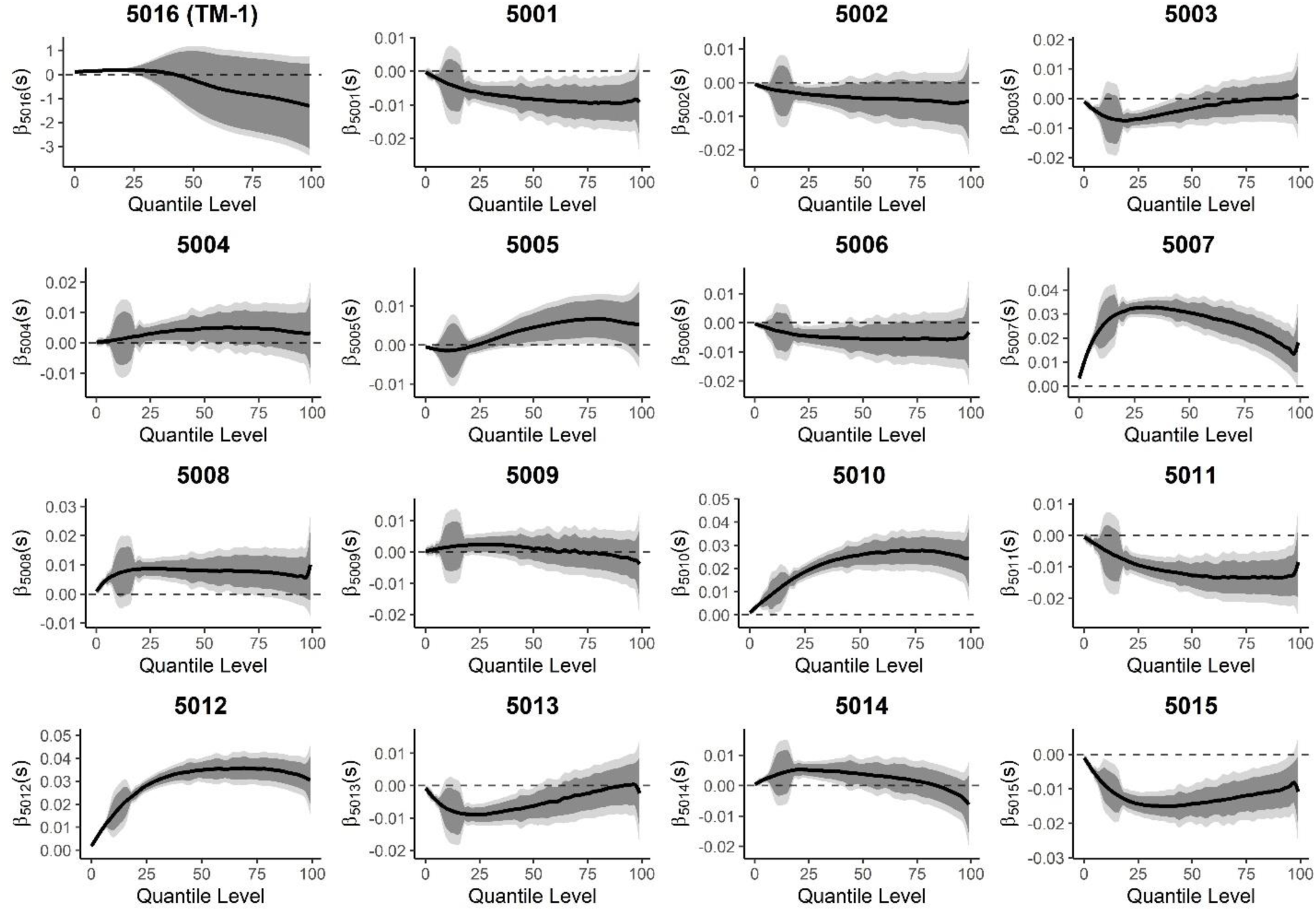
Function-on-scalar regression for 16 different cotton genotypes quantified significance of cotton introgression segments on temporal phenomic data (the ExG2 vegetation index). Eq. 1 tested the hypothesis that unique introgression segments each cotton genotype possesses affected spectral characteristics throughout growth across all days after transplanting (DATs). Pointwise unadjusted confidence intervals (dark gray) are shown within correlation and multiplicity adjusted confidence intervals (light gray) (Crainiceanu et al., 2024). Specific regions of the quantile domain with different Genotype effects are significant at a quantile level(s) if the confidence intervals do not overlap with 0. Genotype 5016 was the reference, and other estimates 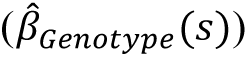 are interpreted as the mean difference in ExG2 values between that genotype and the reference at quantile level 𝑝.

**Table 2.**
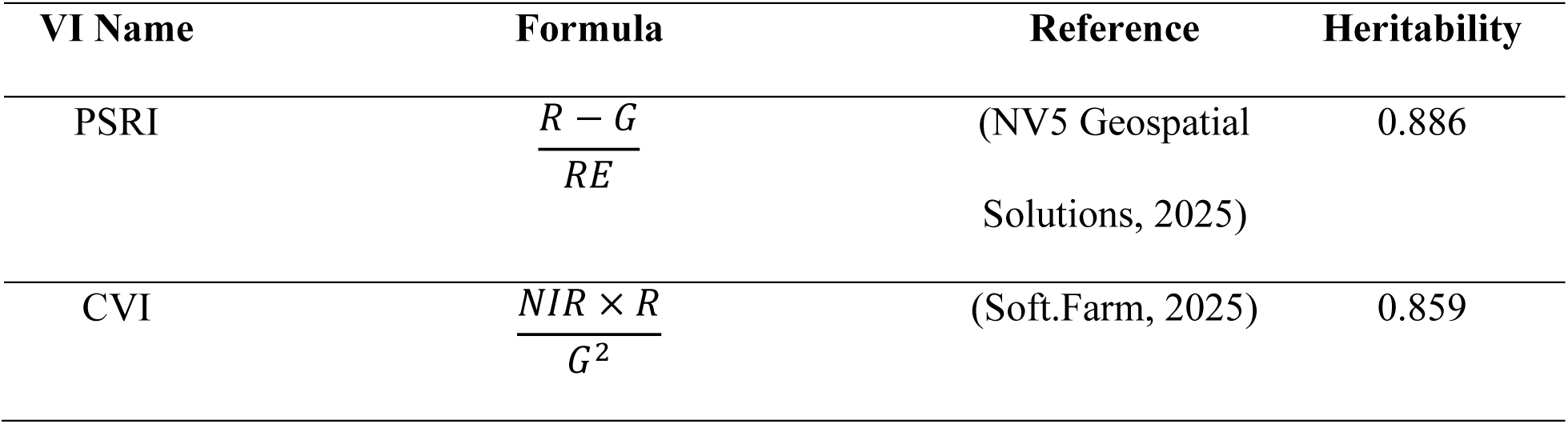
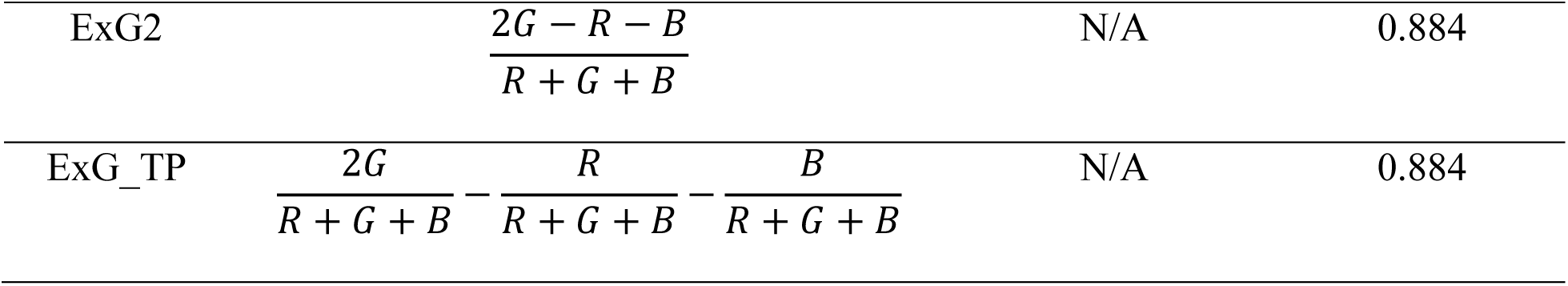
Highest-heritability VI formulas and heritabilities for multispectral and RGB VIs in single plant cotton analysis. Values denote the median across all DATs and quantile levels from HM3.

| VI Name | Formula | Reference | Heritability |
| --- | --- | --- | --- |
| PSRI | $\frac{R - G}{RE}$ | (NV5 Geospatial Solutions, 2025) | 0.886 |
| CVI | $\frac{NIR \times R}{G^2}$ | (Soft.Farm, 2025) | 0.859 |
| ExG2 | $\frac{2G - R - B}{R + G + B}$ | N/A | 0.884 |
| ExG_TP | $\frac{2G}{R + G + B} - \frac{R}{R + G + B} - \frac{B}{R + G + B}$ | N/A | 0.884 |

### Temporal Drone Flights Provide Whole-Season Insights into Heritability and Variance Components for Cotton and Maize

Daily drone flights (cotton) and sparse temporal G2F flights (maize) quantified the influence of Genotype on routine (median values per time point) phenomic data when analyzed at the whole-season level (AM1) and for each flight (AM2) with median VI data (Supplementary Results; Supplementary Fig. 1; Fig. S1).

### Novel Quantile Levels are more Heritable than the Median and Show Dynamic Within-Day Variance Components and Heritability

Within-day variance decomposition was explored to assess heritability and explanatory power of Genotype for VI data along the quantile domain for each flight date. A large volume of quantitative tabular data was produced per cotton plant (989,800 points per plant; 98 VIs × 101 flight dates × 100 quantile levels) and maize plot (62,900 data points per plot; 37 VIs × 17 flight dates × 100 quantile levels in TXH1.2020), 100-fold more than the mean or median alone. For cotton, the median tended not to be the most heritable quantile level regardless of the VI analyzed (Table 3). Similar results were observed with aggregated data from all 11 G2F environments, with an exception noted in TXH1.2020 (Tables S2-3).

**Table 3.**
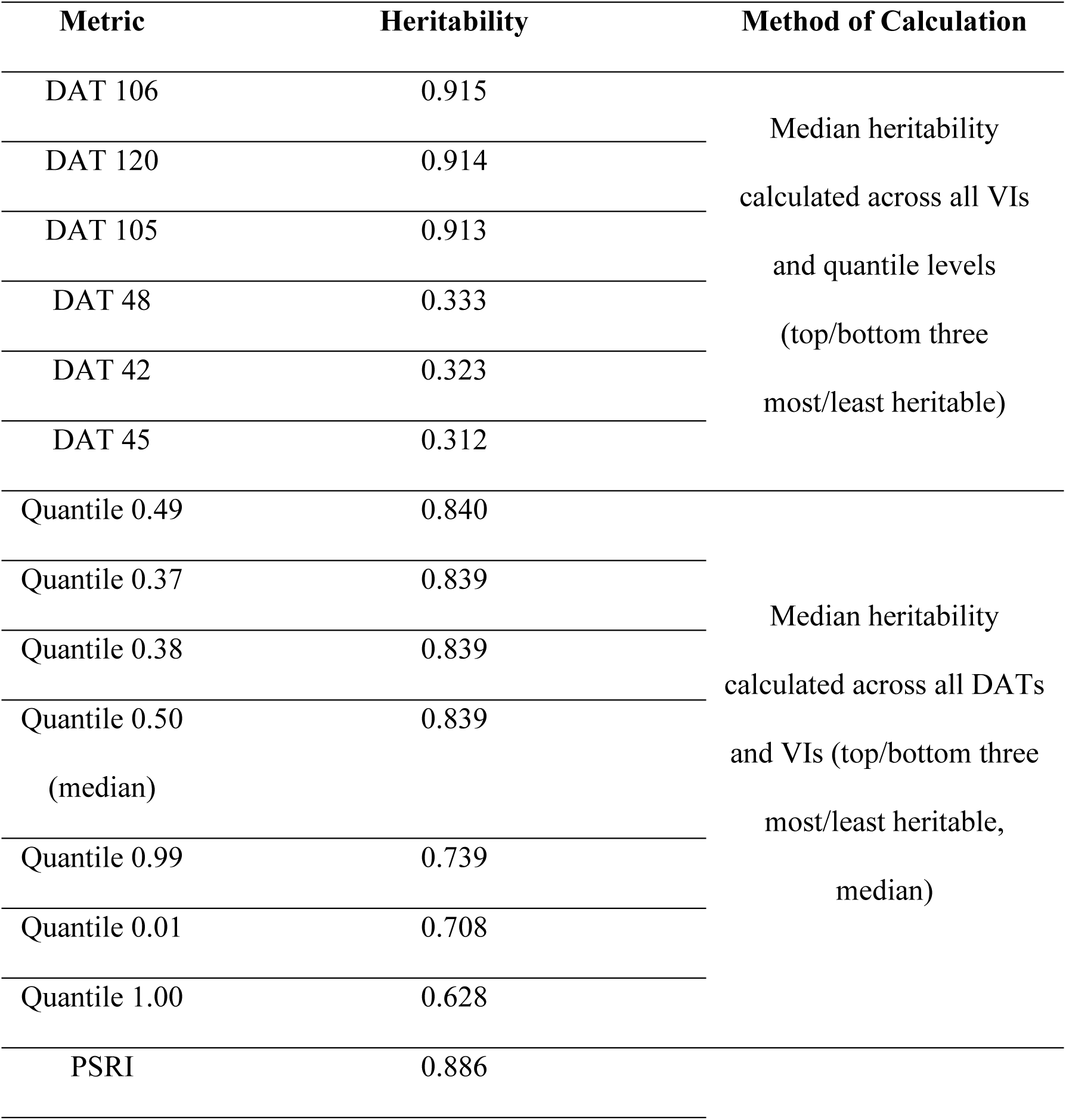
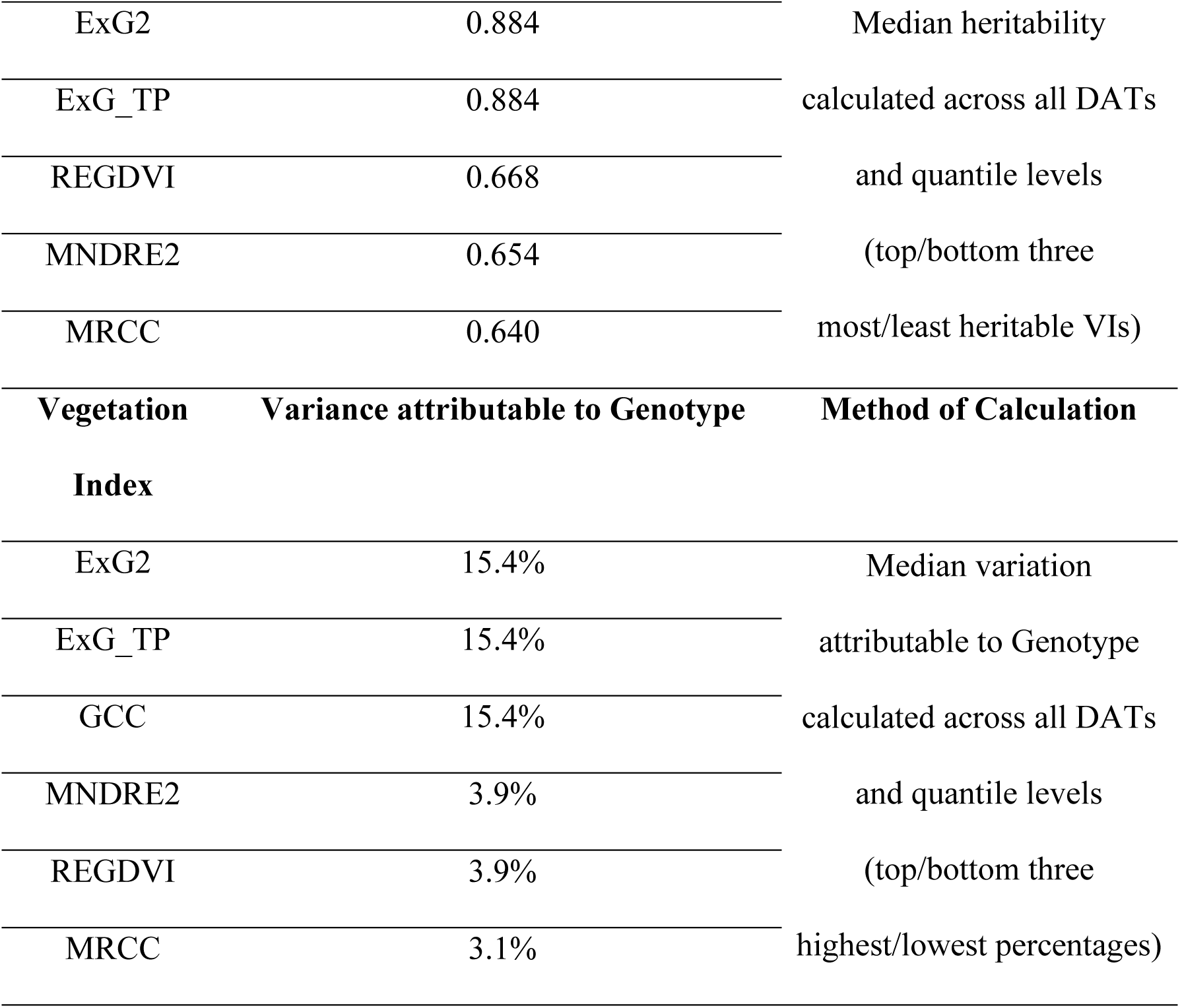
Range of highest and lowest heritability days, VIs, and quantile levels in single plant cotton model HM3 along with genetic variance. The median was used to summarize across all other metrics than those presented. In each case, the top/bottom three most/least heritable results are presented in descending order.

*Cotton*: the median was the 46^th^ most heritable quantile level (𝐻^2^= 0.916) across five example DATs and four VIs (Fig. 4A) and 13^th^ across all dates (𝐻^2^= 0.839).These models extend mixed models to be applied at each quantile level separately within each DAT, facilitating variance component and heritability analysis across the quantile functional domain and the time domain. Heritability tapered at the extremes of the quantile domain (toward 0.01 and 1.00, Fig. 4A), which was expected given these regions capture outlier plant pixel values in each VI distribution for each plant. Five days throughout growth were chosen across the season to highlight differences in variance in the VIs and heritability attributable to Genotype, both across the time and quantile domains. All cotton VIs showed trends of increasing Genotype variance toward the middle of the season, consistent with standard VIs (Fig. S1B), while maize Genotype variance was highest toward the end of the season (Fig. 4B; Figs. S4-14). More genetic variance was observed in maize than cotton because the G2F maize population was larger and more diverse while the cotton population was small (16 genotypes) and nearly isogenic. Heritability was higher in cotton because there were more replicates that minimized the influence of residual error (HM3).

**Figure 4.**
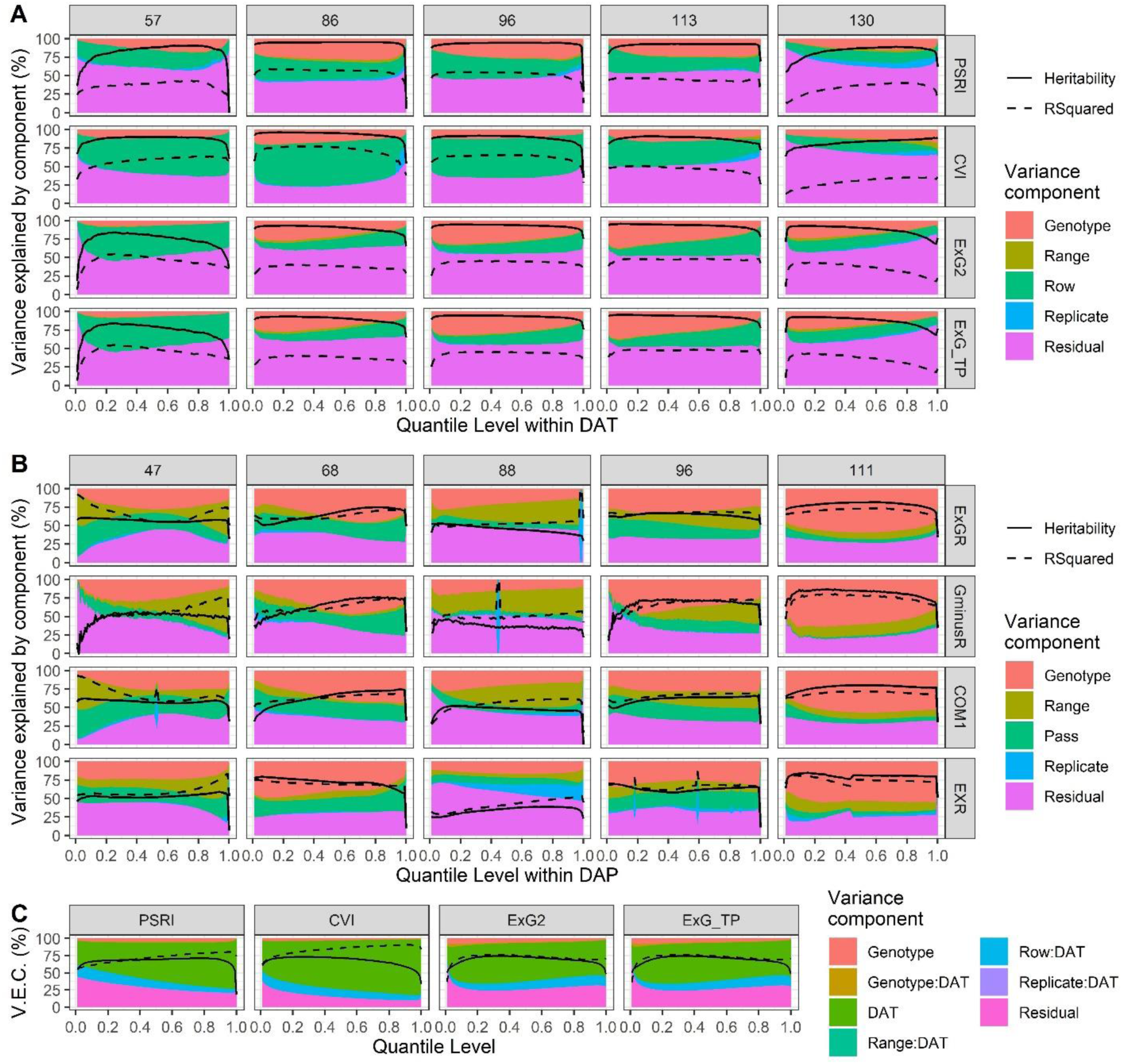
Variance explained by component (V.E.C.) for different quantile levels within (**A**-**B**) and across (**C**) time points in cotton and maize. (**A**) Variance components (AM3) and heritability (HM3) in cotton across all quantiles for five example dates. An ANOVA model (AM3) was evaluated for each combination of vegetation index (VI), day after transplanting (DAT), and quantile level, producing the most granular analysis tested (quantiles 0.01-1.00 in increments of 0.01 per DAT). (**B**) Similar model results for TXH1.2020 G2F maize data; DAP – days after planting; (**C**) Variance components (AM4) and heritability (HM4) after analyzing the distributional vegetation index (VI) data set (cotton) within each quantile level across time points (days after transplanting, DAT). AM4 and HM4 evaluated Genotype × DAT interaction across the quantile domain. VI acronyms: PSRI – plant senescence reflectance index; CVI – chlorophyll VI; ExG2 – modified excess green VI; ExG_TP – modified excess green VI; COM1 – combined index 1; ExGR – modified excess green VI; ExR – excess red VI; GminusR – green minus red index.

Fluctuations of heritability were observed from day to day, so the only recorded confounding variable, the hour of flights, was investigated. AM2 revealed nonsignificant correlations between flight time of day and heritability in both cotton and maize (Figs. S2-3), indicating detectable changes in developmental processes in adjacent flights not attributable to imaging and lighting effects.

*Maize*: when data were pooled across all 11 G2F environments from AM3 variance components, the median was the 14^th^ most heritable quantile level (𝐻^2^= 0.633). However, across 17 flights in one location, TXH1.2020 (100 quantile levels × 17 flights = 1,700 ANOVAs), the median was the most heritable quantile level (𝐻^2^ = 0.563) aggregated for all flight dates and VIs (Table S2). The VI explained best by Genotype in TXH1.2020 was ExGR (28.6%; Table S2) across all flight dates and quantile levels, which was also true when aggregating data from 11 environments (ExGR Genotype variance = 32.2%; Table S3). The most heritable VIs and their formulas in TXH1.2020 and aggregated across 11 G2F environments are given in tables S4 and S5, respectively.

### Temporal Distributional Data Enabled Analysis of Genotype×Time Interaction at Different Quantiles (AM4 and HM4) and Genotype×Quantile Interaction at Different Days (AM5 and HM5)

AM4 and HM4, testing for Genotype × Time interactions, revealed the 99 other ANOVA and heritability results (Fig. 4C) not possible to calculate with AM1 and HM1 that only considered the median of the VI distributions (quantile level 0.50). Similarly, AM5 and HM5 enabled quantification of Genotype × Quantile interactions at each step along the quantile domain (Fig. S15). RGB VIs (ExG2 and ExG_TP) were best explained by Genotype in the lower regions of the quantile domain using AM4 with the added Genotype × DAT interaction effect not included in AM3 (Fig. 4C), suggesting this phenomenon was stable across time in this experiment.

### Heritability Attention Maps (HAMs) Link Per-Pixel Heritability with Developmental Stages

Heritability attention maps (HAMs) generated from Eq. 2 for cotton (Fig. 5A) and maize (Fig. 5B) revealed the pixel-by-pixel genetic nature of the distributional data contained in each image of a plant (cotton) or across a plot (maize). These visualizations provide a granular and explainable depiction of the mixed model results (Fig. 4A-B), highlighting specific regions of plants and plots with high or low heritability. The highest heritability was observed close to 90 DAT in cotton, with less heritable pixel values in early and late flights. For maize, heritability generally increased toward the end of the season (e.g., 111 DAP), yet higher heritability flights were also observed during tasseling (e.g., 68 DAP), and vegetative growth (e.g., 47 DAP). Lower heritability dates (e.g., 88 DAP) interrupted high heritability dates. Given the nonsignificant correlations between heritability and flight time of day in both cotton and maize (Figs. S2-3) this was most likely an unknown biological phenomenon.

**Figure 5.**
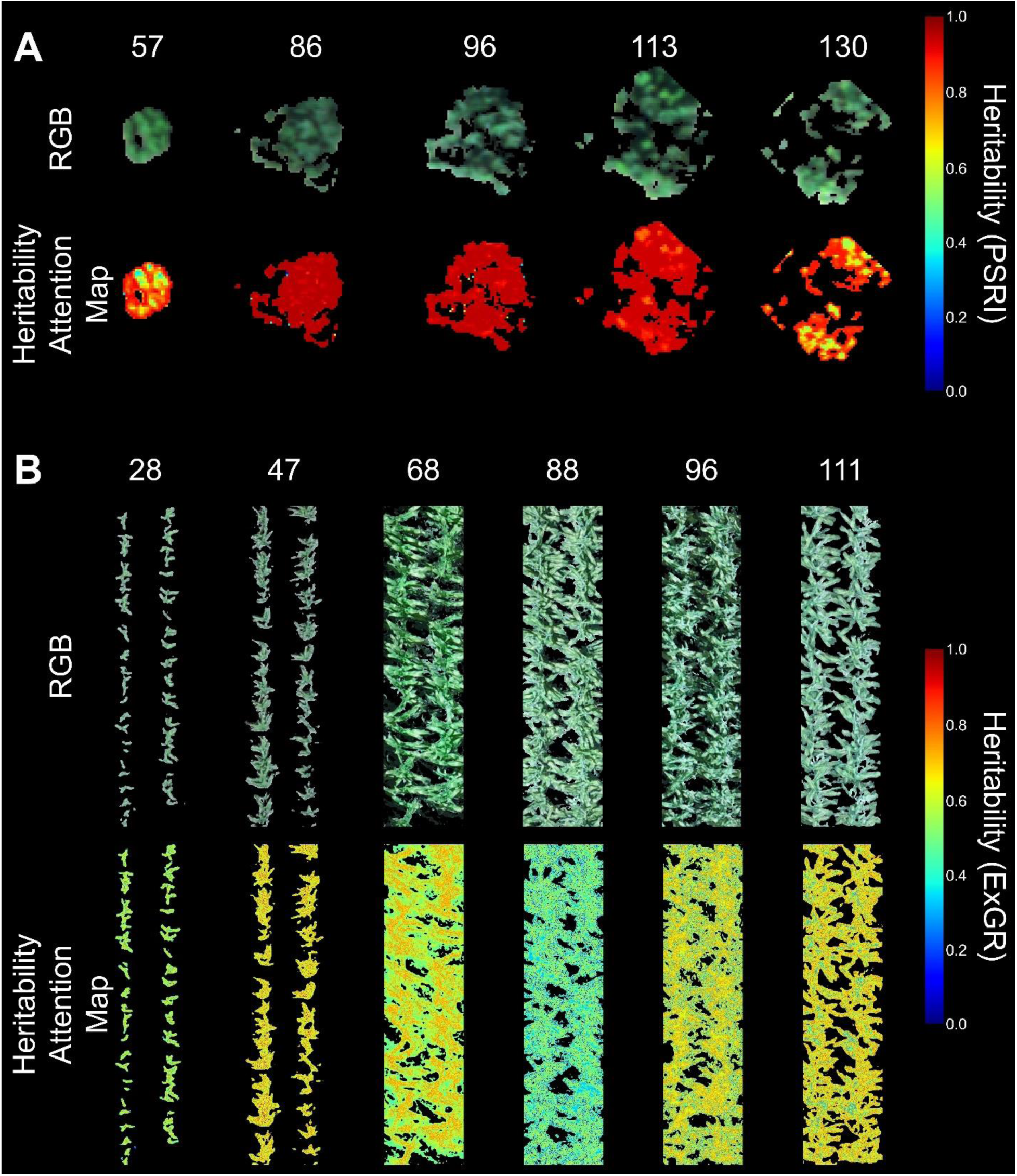
Heritability attention maps (HAMs) for cotton (**A**) and maize (**B**). Heritability data was used from AM3 in Eq. 2 with all RGB images associated with each time point. Per-pixel heritability values are given for single-plant cotton images where numbers at the top of each image/HAM pair indicate the flight date in days after transplanting (DAT) for cotton (**A**) or planting (DAP) for TXH1.2020 maize (**B**). Maize day 88, with low heritability, was chosen in contrast to flanking higher heritability dates.

### Distributional Data Enable Higher Grain Yield Predictions than the Median Alone in 11 G2F Environments

Elastic-net regression enabled 1) direct comparisons of prediction ability between median-only and distributional predictors (Fig. 6) and 2) quantification of variable importance scores for predicting grain yield from distributional data across 11 G2F environments (TXH1.2020 in Fig. 7; other environments in Supplementary Figs. 16-25). Across and within every G2F environment, at least one model trained with distributional data predicted yield with higher correlations to actual yield than median-only regressions (Fig. 6). Aggregated across environments, distributional predictor regression outperformed median regression by 12.7%-21.6% (Fig. 6, top left). The extreme distributional quantiles were less heritable (Fig. 4), containing spurious points and pixels. Descending from the top of Fig. 7 are stricter levels of data quantile truncation, starting with retaining all data, removing quantiles 0.01 and 1.00, and lastly removing quantiles 0.01-0.25 and 0.76-1.00. Surprisingly, distributional predictors other than the median were retained by shrinkage and variable selection. For example, in TXH1.2020 DAP 68 (tassel emergence in this environment), the 0.39 quantile level had nonzero variable importance scores in all data filtration scenarios (Supplementary Fig. 26; Videos S2-4). Similar phenomena were observed in other environments such as WIH2.2021 DAP 111, where the 0.28 quantile level had variable importance scores ranging between 10.1% and 35.3% for all scenarios (Fig. S23).

**Figure 6.**
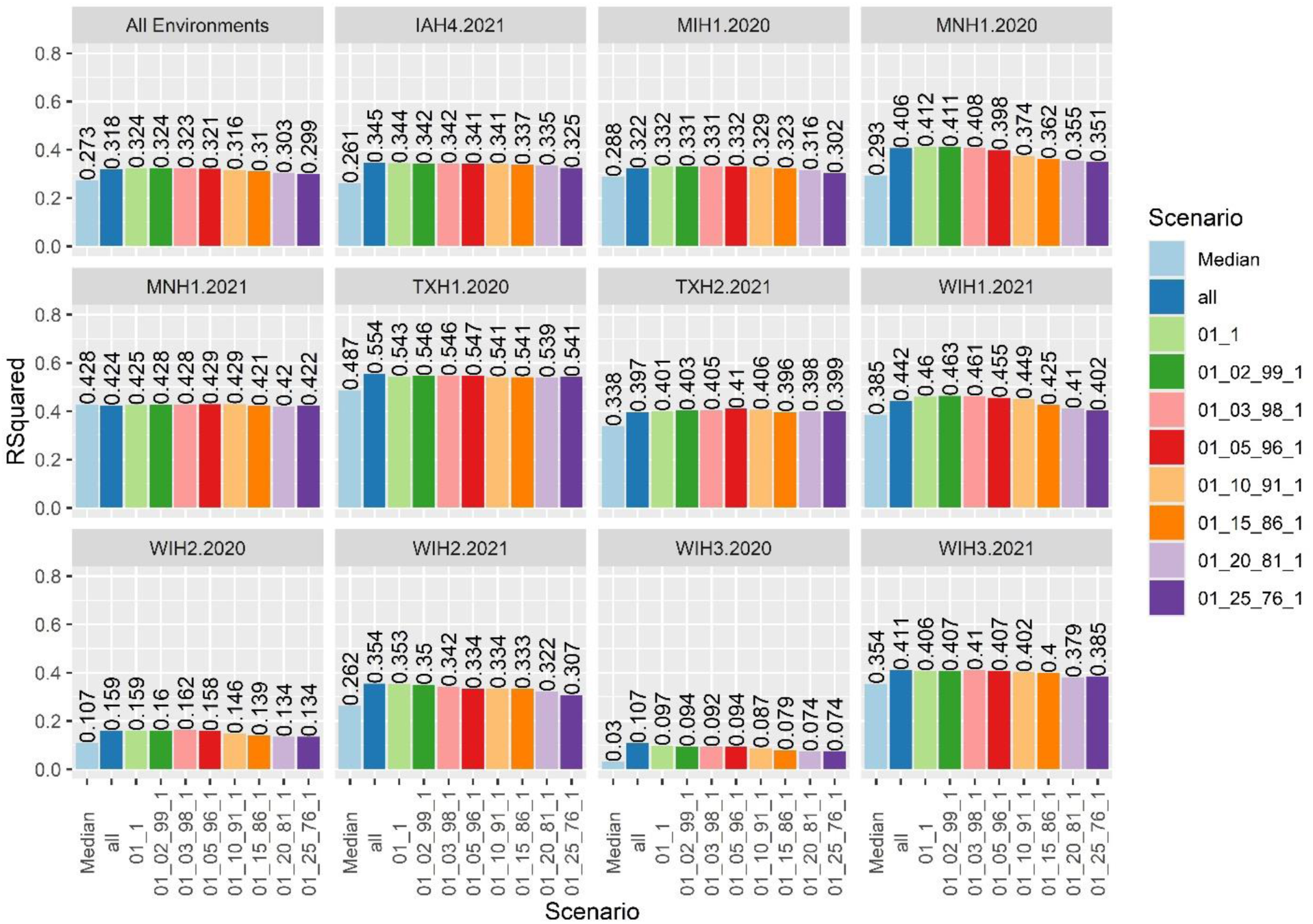
Correlations between actual and predicted grain yield values after training elastic-net regressions with ExGR vegetation index (VI) data to predict grain yield separately within each environment. The “All Environments” pane reports average correlations across all the G2F environments in the analysis.“Median” indicates that regressions had access to only the median VI values; “all” indicates all data were retained as predictors (100 quantile levels × the number of flights), “01_1” indicates quantile levels 0.01 and 1.00 were removed, and “01_25_76_1” indicates quantile levels between 0.01 to 0.25 and between 0.76 to 1.00 were removed before training and testing the elastic-net regression.

**Figure 7.**
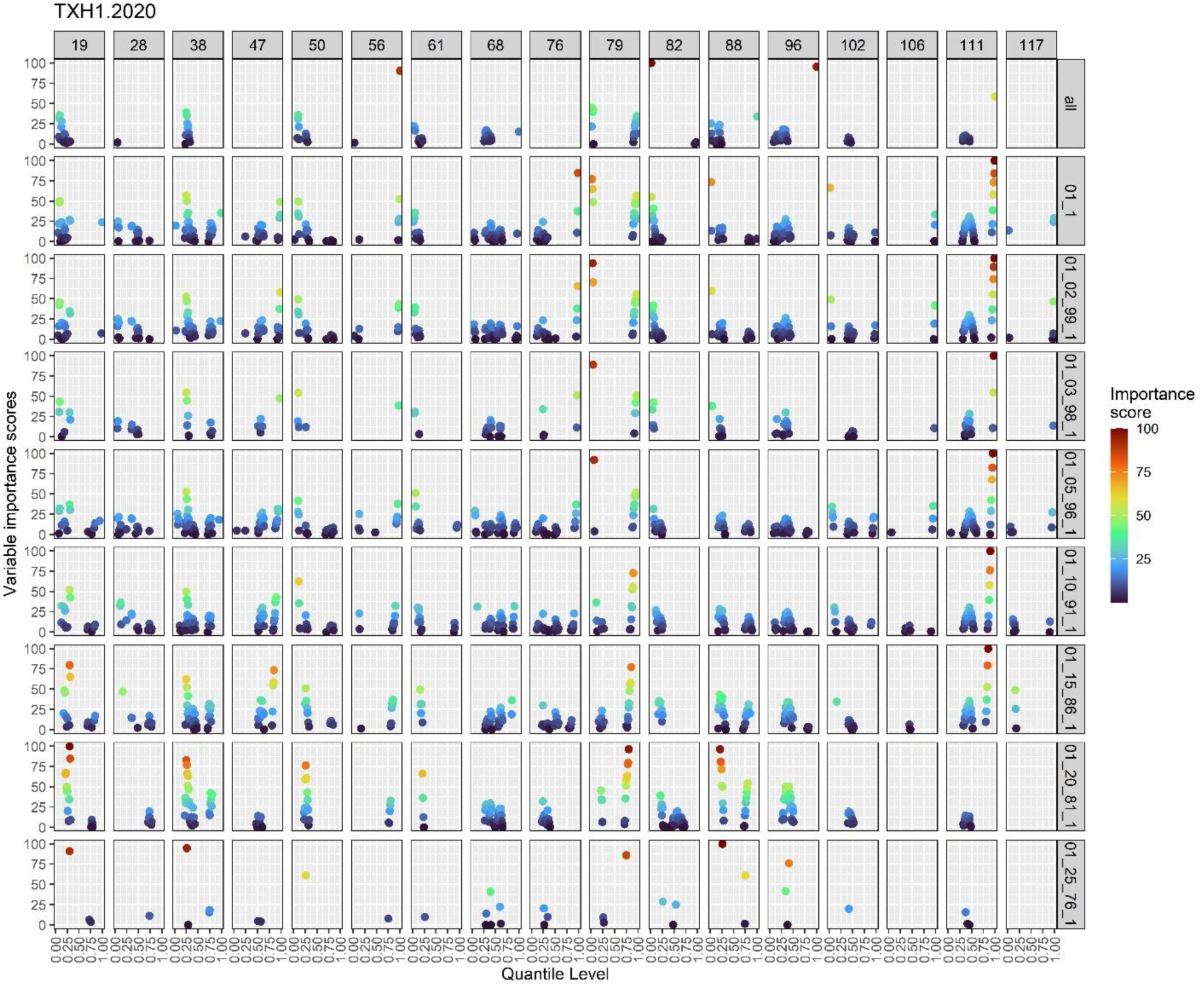
Variable importance scores using ExGR distributional data as predictors of grain yield in TXH1.2020 G2F maize. Column titles are drone flight dates in days after planting (DAP). Row labels indicate the level of data filtration of the distributional predictors. “All” indicates that all data were kept as predictors (100 quantile levels × the number of flight dates), “01_1” indicates quantile levels 0.01 and 1.00 were removed, with “01_25_76_1” indicating that quantile levels between 0.01 to 0.25 and between 0.76 to 1.00 were removed before training and testing the elastic-net regression. A zoomed in image of day 68 (Fig. S26) shows clumps of points are relatively smooth curves. Videos S2-4 show the progression of regions of three maize plots highlighted by each quantile level.

### Correlation Gradients of Temporal Quantile Phenomic Features

A correlation analysis was conducted to screen for redundancy of distributional data.

Fisher-adjusted mean correlations among quantile levels across DAPs within G2F environments and pooled across all 11 environments revealed decaying temporal correlations moving from one side of the quantile domain to the other (Fig. 8). This demonstrated that adjacent quantiles (e.g., 0.25 and 0.26) were highly correlated, but distant quantiles (e.g., 0.25 and 0.75) added unique information as seen in the elastic-net prediction variable importance results (Fig. 7; Figs. S16-26). The environments with the narrowest and widest ranges of correlation coefficients were MIH1.2020 (0.52-1.00) and MNH1.2020 (0.09-1.00), respectively, while correlations ranged between 0.25-1.00 when data were combined across all environments.

**Figure 8.**
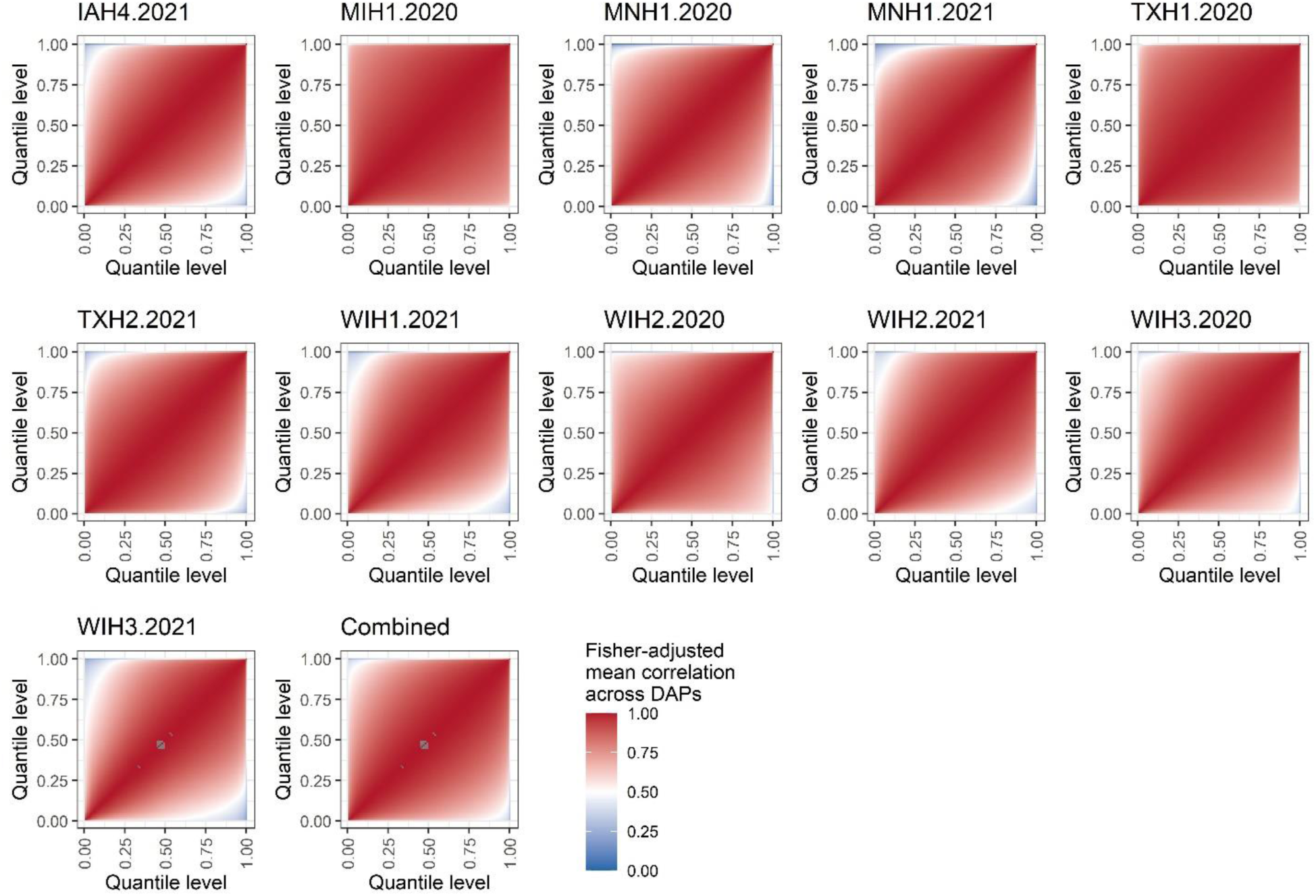
Fisher-adjusted mean correlations were calculated across days after planting (DAPs) for each maize G2F environment separately and across DAPs for data pooled across all 11 environments (the “Combined” panel of the figure). Temporal correlations between quantile levels decayed moving from one side of the quantile domain to the other.

## Discussion

Overall, the analyses presented for cotton and maize demonstrated that distributional quantile data captured novel temporal dynamics of plant growth and senescence through three case studies. i) A novel mixed function-on-scalar distributional regression implementation for cotton phenomic data revealed statistically significant effects of inter-species introgression segments at specific regions of the quantile domain. ii) Elastic-net regression showed that distributional maize data was both useful for prediction ability, leading to higher correlations between actual and predicted yield, and important, as shown by nonzero elastic-net variable importance scores at quantiles besides the median. iii) Within-day variance decomposition was supported by a novel per-pixel heritability visualization method that revealed developmental stage-specific explainability (e.g., high 𝐻^2^ pixels near male anthers during flowering in maize and toward the edges of leaves at the onset of senescence in both crops). Python scripts developed for this study can be extended to other crops and potentially other kingdoms’ temporal imaging data. The explainability of the proposed methods coupled with application across two crops and multiple environments indicate the importance of distributional data analysis for discovering developmental biological features, prediction of end traits, and characterizing phenotypic plasticity, with applications for understanding plant responses to abiotic stresses.

Several challenges of using phenomic features from temporal imaging for biological discovery and for predicting end traits across diverse environments were addressed. One challenge is a lack of numerous, consistent, annotated phenomic feature predictors (like gene ontologies), addressed by extracting different quantiles within experimental units at each time point. This distributional data approach assessed 100× more features than were previously available through quantiles in 0.01 increments for each image. Although adjacent quantiles were highly correlated, lower correlations (Fig. 8) and importance in yield predictions were observed with distanced quantiles (see quantile clusters in Fig. 7; Fig. S26). Another challenge is in integrating developmentally unlinked observations and sparse temporal data across environments. In the maize G2F case, populations experienced different weather and photoperiods, and thus biological development across each environment, making DAP technically but not biologically optimal for alignment. Even within environments, genotypes can perceive photoperiods differently conditioned on temperature (Drouault et al., 2025). Furthermore, for maize, each environment’s images were sparsely collected at different times. Finally, lack of explainability (common with deep learning frameworks) is a major challenge of high-dimensional phenomic data, which cannot be manually measured or “ground-truthed”. This was addressed by developing heritability attention maps (HAMs), providing a pixel-by-pixel overlay of the heritability of image features for each VI within each observation date. HAMs revealed that among the most heritable quantile features in maize were pixels around the tassels at flowering (DAP 68, Fig. 5B) as well as the leaf edges at the initiation of senescence (DAP 111). While deep learning methods continue to demonstrate consistently impressive results for plant imaging applications, they suffer from varying degrees of explainability (Mostafa et al., 2023; Rudin, 2019) that complicates direct applicability. Explainability was facilitated through HAMs coupled with function-on-scalar regression that visualized and highlighted the location in the continuum (quantile in this case, though time could be used) and magnitude of the influence of predictors such as Genotype (Fig. 3).

Viewing plant phenomic data as distributions across the range of image pixels instead of single number summaries per image (e.g., mean, median) was motivated by analyses of medical wearable data (Ghosal & Matabuena, 2024; Matabuena & Crainiceanu, 2024). Visualization allowed novel insights to be derived from distributional data, such as a unique trajectory from planting to peak greenness through senescence for the individual plant or plot chosen for the visualization (shape of the sheet in Fig. 2). Such insights are lost by only retaining scalar summary trajectories for downstream analysis. While VIs were used in this study, the methods are extensible to other phenomic features currently described by summary statistics, such as texture and canopy structural variation, which could lead to distributional data calculated from 3D models of experimental units instead of the 2D images used in this study.

Temporal quantile data afforded multiple types of mixed model analyses (Table 1), the most granular of which was AM3 and its associated heritability model (HM3). Variance components and the resulting heritabilities in cotton (Fig. 4A) and maize (Fig 4B; Figs. S4-14) were dynamic both within and across drone flight dates (Fig. 4C). Traditional VI scalar summaries were not the most heritable for data pooled across 11 sets of maize G2F flights or across the 101 cotton flights. Interestingly, variance attributable to cotton Genotype and resulting heritability when analyzing only the median quantile (Fig. S1B) tended to be highest toward the middle of the season. However, this median-only analysis missed quantile levels that exceeded the median heritability both in cotton (Fig. 4A) and maize (Fig. 4B). Distributional data analysis has the potential to provide richer insights into plant development and to improve trait prediction (Fig. 6), all without requiring any modifications to existing phenomic data capture protocols.

Further, sensitivity analyses of elastic-net models across 11 maize G2F environments demonstrated that distributional predictors explain new biological information not captured by the mean or median, indicating non-median quantile levels consistently had non-zero (often substantially above zero) variable importance scores even as quantiles at the edges of the distribution were removed (Fig. 7). Grain yield prediction ability improvements of elastic-net regressions trained with distributional predictors vs. median-only predictors ranged between 12.7%-21.6%. In TXH1.2020, the 0.39 quantile level had variable importance scores ranging between 10.0% and 40.9% across nine data filtration levels (Fig. S26). Intriguingly, dates with minimal genetic signal occasionally appeared predictive, presumably due to leverage from residual soil/background pixels, thus motivating stricter plant-soil segmentation (e.g., Otsu (Otsu, 1975)) (Fig. 7; Figs. S16-25, Videos S2-4).

The results from the fast univariate implementation of the functional mixed model (Eq. 1) (Cui et al., 2022; Loewinger et al., 2025) allowed interpreting the effects of cotton Genotypes on specific regions of the quantile domain. This novel model tested the hypothesis that different genomic introgression regions exerted a significant effect on phenomic data. Other fixed effects covariates of interest could be included, and *lme4* users will recognize an intuitive approach to programming fixed and random effects in the *fastFMM* package (Loewinger et al., 2025). The phenotypic stability of cotton lines evaluated in this study with their prior performance (DeSalvio et al., 2024) suggests the presence of *G. tomentosum* introgressions that could be targeted for drought-or stress-tolerance breeding.

## Conclusions

Considering pixels from temporal images of individual experimental units as distributions instead of scalar summaries can provide novel biological insights and enhance prediction abilities. Analyses of quantile functions evaluated for images of single plants and plots discovered more heritable phenomic features than conventional mean and median scalar summary values. Many quantiles were imperfectly correlated with each other and with the median. The novel functional mixed model (FMM) proposed presents the first analysis of functional fixed effects for temporal drone imagery. Sensitivity analyses indicated that despite removing large parts of the quantile domain, stable distributional predictors for maize grain yield could be identified, supporting the improved prediction abilities of 12.7%-21.6% facilitated by distributional predictors over the models trained with the median. Lastly, a novel data visualization method (heritability heatmaps, HAMs) was proposed to link heritability to pixels within images of single experimental units (plants or plots), highlighting biologically relevant signals such as maize tassel emergence and senescence initiation within phenomic distributional quantile data.

## Acknowledgements

The authors would like to acknowledge the dedication and efforts of Dr. Robert Vaughn and the undergraduate students in the Stelly cotton research group (Texas A&M University) for technical support needed to sow, spaced-transplant, grow and maintain the cotton plots, Amanda Gilbert and Dorothy Sweet (field technical support and drone data acquisition; University of Minnesota), Dr. Maninder Singh (planting and harvesting field trial; Michigan State University), Lindsey Newton (field data collection and analysis; Michigan State University), Erin Bunting and Robert Goodwin (drone data acquisition and preprocessing; Michigan State University), Colby Bass (field data collection and field maintenance; Texas A&M University), and the graduate and undergraduate students and high school employees of the Texas A&M Quantitative Genetics and Maize Breeding Program for many hours of phenotypic data collection and field maintenance. Mention of trade names or commercial products in this publication is solely for the purpose of providing specific information and does not imply recommendation or endorsement by the U.S. Department of Agriculture. USDA is an equal opportunity provider and Employer. Any opinions, findings, and conclusions or recommendations expressed in this material are those of the authors(s) and do not necessarily reflect the views of the National Science Foundation.

## Data Availability

All code used in this research is available at the corresponding GitHub repository, along with files needed to reproduce the results: https://github.com/ajdesalvio/phenomic-distributional

## References

Acin-Albiac, M., Filannino, P., Gobbetti, M., & Di Cagno, R. (2020). Microbial high throughput phenomics: The potential of an irreplaceable omics. Computational and Structural Biotechnology Journal, 18, 2290–2299.

Adak, A., Murray, S. C., Božinović, S., Lindsey, R., Nakasagga, S., Chatterjee, S., Anderson, S. L., & Wilde, S. (2021). Temporal Vegetation Indices and Plant Height from Remotely Sensed Imagery Can Predict Grain Yield and Flowering Time Breeding Value in Maize via Machine Learning Regression. Remote Sensing, 13(11). 10.3390/rs13112141

Adunola, P., Tavares Flores, E., Riva-Souza, E. M., Ferrão, M. A. G., Senra, J. F. B., Comério, M., Espindula, M. C., Verdin Filho, A. C., Volpi, P. S., & Fonseca, A. F. A. (2024). A comparison of genomic and phenomic selection methods for yield prediction in Coffea canephora. The Plant Phenome Journal, 7(1), e20109.

Bates, D., Mächler, M., Bolker, B., & Walker, S. (2015). Fitting linear mixed-effects models using lme4. Journal of statistical software, 67, 1–48.

Chatterjee, S., Murray, S. C., Matias, F. I., & Fahlgren, N. (2025). FIELDimagePy: A tool to estimate zonal statistics from an image, bounded by one or multiple polygons. Crop Science, 65(1), e21357.

Crainiceanu, C. M., Goldsmith, J., Leroux, A., & Cui, E. (2024). Functional data analysis with R. CRC Press.

Cui, E., Leroux, A., Smirnova, E., & Crainiceanu, C. M. (2022). Fast univariate inference for longitudinal functional models. Journal of Computational and Graphical Statistics, 31(1), 219–230.

DeSalvio, A. J., Adak, A., Arik, M. A., Shepard, N. R., DeSalvio, S. M., Murray, S. C., García-Ramos, O., Badavath, H., & Stelly, D. M. (2024). Temporal image sandwiches enable link between functional data analysis and deep learning for single-plant cotton senescence. in silico Plants, 6(2), diae019.

Drouault, J., Palaffre, C., Millet, E. J., Rodriguez, J., Martre, P., Johnson, K., Parent, B., Welcker, C., & Wisser, R. J. (2025). A reaction norm for flowering time plasticity reveals physiological footprints of maize adaptation. G3: Genes, Genomes, Genetics, jkaf095.

Fisher, R. A. (1915). Frequency distribution of the values of the correlation coefficient in samples from an indefinitely large population. Biometrika, 10(4), 507–521.

Friedman, J., Hastie, T., Tibshirani, R., Narasimhan, B., Tay, K., Simon, N., & Qian, J. (2021). Package ‘glmnet’. CRAN R Repositary, 595.

Furbank, R. T., & Tester, M. (2011). Phenomics–technologies to relieve the phenotyping bottleneck. Trends in Plant Science, 16(12), 635–644.

Gano, B., Bhadra, S., Vilbig, J. M., Ahmed, N., Sagan, V., & Shakoor, N. (2024). Drone-based imaging sensors, techniques, and applications in plant phenotyping for crop breeding: A comprehensive review. The Plant Phenome Journal, 7(1), e20100.

Ghosal, R., & Matabuena, M. (2024). Multivariate Scalar on Multidimensional Distribution Regression With Application to Modeling the Association Between Physical Activity and Cognitive Functions. Biometrical Journal, 66(7), e202400042.

Ghosal, R., Varma, V. R., Volfson, D., Hillel, I., Urbanek, J., Hausdorff, J. M., Watts, A., & Zipunnikov, V. (2023). Distributional data analysis via quantile functions and its application to modeling digital biomarkers of gait in Alzheimer’s disease. Biostatistics, 24(3), 539–561.

Guo, W., Carroll, M. E., Singh, A., Swetnam, T. L., Merchant, N., Sarkar, S., Singh, A. K., & Ganapathysubramanian, B. (2021). UAS-based plant phenotyping for research and breeding applications. Plant Phenomics.

Holland, J. B., & Piepho, H.-P. (2024). Don’t BLUP Twice. G3: Genes, Genomes, Genetics, 14(12), jkae250.

Hotelling, H. (1953). New light on the correlation coefficient and its transforms. Journal of the Royal Statistical Society. Series B (Methodological), 15(2), 193–232.

Houle, D., Govindaraju, D. R., & Omholt, S. (2010). Phenomics: the next challenge. Nature Reviews Genetics, 11(12), 855–866.

Huang, C., Thompson, P., Wang, Y., Yu, Y., Zhang, J., Kong, D., Colen, R. R., Knickmeyer, R. C., Zhu, H., & Alzheimer’s Disease Neuroimaging, I. (2017). FGWAS: functional genome wide association analysis. Neuroimage, 159, 107–121.

Jackson, R., Buntjer, J. B., Bentley, A. R., Lage, J., Byrne, E., Burt, C., Jack, P., Berry, S., Flatman, E., & Poupard, B. (2023). Phenomic and genomic prediction of yield on multiple locations in winter wheat. Frontiers in genetics, 14, 1164935.

Kaya, C. (2025). Optimizing Crop Production With Plant Phenomics Through High-Throughput Phenotyping and AI in Controlled Environments. Food and Energy Security, 14(1), e70050.

Klein, N. (2024). Distributional regression for data analysis. Annual Review of Statistics and Its Application, 11.

Kuhn, M. (2008). Building predictive models in R using the caret package. Journal of statistical software, 28(1), 1–26.

Kumar, J., Pratap, A., & Kumar, S. (2015). Plant phenomics: an overview. Phenomics in crop plants: trends, options and limitations, 1-10.

Lane, H. M., Murray, S. C., Montesinos-López, O. A., Montesinos-López, A., Crossa, J., Rooney, D. K., Barrero-Farfan, I. D., De La Fuente, G. N., & Morgan, C. L. S. (2020). Phenomic selection and prediction of maize grain yield from near-infrared reflectance spectroscopy of kernels. The Plant Phenome Journal, 3(1), e20002.

Lim, P. O., Kim, H. J., & Gil Nam, H. (2007). Leaf senescence. Annu. Rev. Plant Biol., 58, 115–136.

Lima, D. C., Aviles, A. C., Alpers, R. T., Perkins, A., Schoemaker, D. L., Costa, M., Michel, K. J., Kaeppler, S., Ertl, D., & Romay, M. C. (2023). 2020-2021 field seasons of Maize GxE project within the Genomes to Fields Initiative. BMC research notes, 16(1), 219.

Loewinger, G., Cui, E., Lovinger, D., & Pereira, F. (2025). A statistical framework for analysis of trial-level temporal dynamics in fiber photometry experiments. Elife, 13, RP95802.

Lyu, J. I. L., Baek, S. H., Jung, S., Chu, H., Nam, H. G., Kim, J., & Lim, P. O. (2017). High-throughput and computational study of leaf senescence through a phenomic approach. Frontiers in plant science, 8, 250.

Matabuena, M., & Crainiceanu, C. M. (2024). Multilevel functional distributional models with application to continuous glucose monitoring in diabetes clinical trials. arXiv preprint arXiv:2403.10514v2.

Matabuena, M., Ghosal, R., Aguilar, J. E., Wagner, R., Merino, C. F., Castro, J. S., Zipunnikov, V., Onnela, J.-P., & Gude, F. (2024). Glucodensity functional profiles outperform traditional continuous glucose monitoring metrics. arXiv preprint arXiv:2410.00912.

Matabuena, M., & Petersen, A. (2023). Distributional data analysis of accelerometer data from the NHANES database using nonparametric survey regression models. Journal of the Royal Statistical Society Series C: Applied Statistics, 72(2), 294–313.

Matabuena, M., Petersen, A., Vidal, J. C., & Gude, F. (2021). Glucodensities: A new representation of glucose profiles using distributional data analysis. Statistical methods in medical research, 30(6), 1445–1464.

Matias, F. I., Caraza-Harter, M. V., & Endelman, J. B. (2020). FIELDimageR: an R package to analyze orthomosaic images from agricultural field trials. The Plant Phenome Journal, 3(1), e20005.

Mueller-Sim, T., Jenkins, M., Abel, J., & Kantor, G. (2017, 2017). The Robotanist: A ground-based agricultural robot for high-throughput crop phenotyping.

Muldoon, J. F., & Daynard, T. B. (1981). Effects of within-row plant uniformity on grain yield of maize. Canadian Journal of Plant Science, 61(4), 887–894.

NV5 Geospatial Solutions. (2025). Vegetation Analysis: Using Vegetation Indices in ENVI. NV5. Retrieved 10 April 2025 from https://www.nv5geospatialsoftware.com/Support/Maintenance-Detail/vegetation-analysis-using-vegetation-indices-in-envi

Onnela, J.-P. (2021). Opportunities and challenges in the collection and analysis of digital phenotyping data. Neuropsychopharmacology, 46(1), 45–54.

Otsu, N. (1975). A threshold selection method from gray-level histograms. Automatica, 11(285-296), 23–27.

Parmley, K., Nagasubramanian, K., Sarkar, S., Ganapathysubramanian, B., & Singh, A. K. (2019). Development of optimized phenomic predictors for efficient plant breeding decisions using phenomic-assisted selection in soybean. Plant Phenomics, 2019.

Pawar, P. S., & Matias, F. I. (2023). FIELDimageR. Extra: Advancing user experience and computational efficiency for analysis of orthomosaic from agricultural field trials. The Plant Phenome Journal, 6(1), e20083.

Pérez-Enciso, M., & Steibel, J. P. (2021). Phenomes: the current frontier in animal breeding. Genetics Selection Evolution, 53(1), 22.

Ramsay, J. O., & Silverman, B. W. (2005). Functional data analysis. Springer.

Rincent, R., Charpentier, J.-P., Faivre-Rampant, P., Paux, E., Le Gouis, J., Bastien, C., & Segura, V. (2018). Phenomic selection is a low-cost and high-throughput method based on indirect predictions: proof of concept on wheat and poplar. G3: *Genes, Genomes, Genetics*, *8*(12), 3961-3972.

Robert, P., Auzanneau, J., Goudemand, E., Oury, F.-X., Rolland, B., Heumez, E., Bouchet, S., Le Gouis, J., & Rincent, R. (2022). Phenomic selection in wheat breeding: identification and optimisation of factors influencing prediction accuracy and comparison to genomic selection. Theoretical and Applied Genetics, 1-20.

Soft.Farm. (2025). Vegetation indices NDVI, EVI, GNDVI, CVI, True color. Soft.Farm. Retrieved 10 April 2025 from https://www.soft.farm/en/blog/vegetation-indices-ndvi-evi-gndvi-cvi-true-color-140

Tardieu, F., Cabrera-Bosquet, L., Pridmore, T., & Bennett, M. (2017). Plant phenomics, from sensors to knowledge. Current Biology, 27(15), R770–R783.

Tian, T. S. (2010). Functional data analysis in brain imaging studies. Frontiers in psychology, 1, 35.

Tiberi, S., Crowell, H. L., Samartsidis, P., Weber, L. M., & Robinson, M. D. (2023). distinct: A novel approach to differential distribution analyses. The Annals of Applied Statistics, 17(2), 1681–1700.

Underwood, J., Wendel, A., Schofield, B., McMurray, L., & Kimber, R. (2017). Efficient in-field plant phenomics for row-crops with an autonomous ground vehicle. Journal of field robotics, 34(6), 1061–1083.

Virlet, N., Sabermanesh, K., Sadeghi-Tehran, P., & Hawkesford, M. J. (2016). Field Scanalyzer: An automated robotic field phenotyping platform for detailed crop monitoring. Functional plant biology, 44(1), 143–153.

Waldmann, E. (2018). Quantile regression: A short story on how and why. Statistical Modelling, 18(3-4), 203–218.

Wang, J.-L., Chiou, J.-M., & Müller, H.-G. (2016). Functional data analysis. Annual Review of Statistics and Its Application, 3, 257–295.

Wang, J., Wong, R. K. W., Zhang, X., & Chan, K. C. G. (2023). Flexible functional treatment effect estimation. arXiv preprint arXiv:2309.08039.

White, J. W., Andrade-Sanchez, P., Gore, M. A., Bronson, K. F., Coffelt, T. A., Conley, M. M., Feldmann, K. A., French, A. N., Heun, J. T., & Hunsaker, D. J. (2012). Field-based phenomics for plant genetics research. Field Crops Research, 133, 101–112.

Winn, Z. J., Amsberry, A. L., Haley, S. D., DeWitt, N. D., & Mason, R. E. (2023). Phenomic versus genomic prediction—A comparison of prediction accuracies for grain yield in hard winter wheat lines. The Plant Phenome Journal, 6(1), e20084.

Xue, X., Du, S., Jiao, F., Xi, M., Wang, A., Xu, H., Jiao, Q., Zhang, X., Jiang, H., & Chen, J. (2021). The regulatory network behind maize seed germination: Effects of temperature, water, phytohormones, and nutrients. The Crop Journal, 9(4), 718–724.

Yang, W., Feng, H., Zhang, X., Zhang, J., Doonan, J. H., Batchelor, W. D., Xiong, L., & Yan, J. (2020). Crop Phenomics and High-Throughput Phenotyping: Past Decades, Current Challenges, and Future Perspectives. Molecular Plant, 13(2), 187–214. 10.1016/j.molp.2020.01.008

Ying, W. (2023). Phenomic studies on diseases: potential and challenges. Phenomics, 3(3), 285–299.

Zavafer, A., Bates, H., Mancilla, C., & Ralph, P. J. (2023). Phenomics: conceptualization and importance for plant physiology. Trends in Plant Science, 28(9), 1004–1013.

Zhang, C., McGee, R. J., Vandemark, G. J., & Sankaran, S. (2021). Crop performance evaluation of chickpea and dry pea breeding lines across seasons and locations using phenomics data. Frontiers in plant science, 12, 640259.

Zhao, C., Zhang, Y., Du, J., Guo, X., Wen, W., Gu, S., Wang, J., & Fan, J. (2019). Crop phenomics: current status and perspectives. Frontiers in plant science, 10, 714.

Zhu, X., Leiser, W. L., Hahn, V., & Würschum, T. (2021). Phenomic selection is competitive with genomic selection for breeding of complex traits. The Plant Phenome Journal, 4(1), e20027.

Zou, H., & Hastie, T. (2005). Regularization and variable selection via the elastic net. Journal of the royal statistical society: series B (statistical methodology), 67(2), 301–320.

